# The Impact of Exogenous Mutagens on Human Mitochondrial DNA Ploidy: Analysis of Changes and Possible Protective Mechanisms

**DOI:** 10.1101/2025.05.09.648561

**Authors:** Nikita Van Leiden, Nadezhda Potapova, Natalia Ree

## Abstract

Mitochondrial ploidy — the relative copy number of mitochondrial DNA per mitochondrion — may serve as an early marker of cellular stress and a potential signal of genotoxic damage. In this study, we investigate the dynamics of mitochondrial ploidy in iPSC cells *in vitro* following exposure to a range of chemical mutagens. Using a qPCR-based method allowing accurate quantification of mitochondrial DNA and mitochondria number, we reveal distinct changes in mitochondrial ploidy that correlate with the genotoxic impact of specific agents. Among the mutagens tested, MX, benzidine, cyclophosphamide, hydrogen peroxide, semustine and nickel (II) chloride induced the most pronounced alterations in mitochondrial DNA content and organization. These findings suggest that mitochondrial ploidy can be used as a sensitive molecular indicator of mutagenic stress, potentially reflecting mitochondrial genome maintenance and organelle adaptation mechanisms.

## Introduction

Mitochondria are essential organelles responsible for maintaining cellular energy homeostasis through oxidative phosphorylation, fatty acid metabolism, apoptosis regulation, and intracellular signaling (Spinelli & Haigis, 2018). The human mitochondrial genome (mtDNA), a circular molecule of approximately 16.5 kb, encodes 37 genes necessary for the function of the respiratory chain (Anderson et al., 1981). Unlike the diploid nuclear genome, mtDNA exists in multiple copies per cell — ranging from hundreds to tens of thousands depending on cell type, energy demand, and metabolic state (Robin & Wong, 1988; Wai et al., 2010). This parameter, known as mtDNA ploidy, is a critical determinant of mitochondrial functionality.

Maintaining optimal mtDNA ploidy is essential for adequate gene expression, synthesis of respiratory chain proteins, and adaptation to bioenergetic stress. Both increases and decreases in mtDNA copy number can significantly influence ATP production, the assembly of mitochondrial ribosomes, and the balance of reactive oxygen species (ROS), impacting cell survival and initiating stress response pathways (Gitschlag et al., 2020). Even moderate reductions in copy number can lead to global transcriptional changes in nuclear genes involved in metabolism and stress responses (Finck et.al., 2006, Picard et. al., 2014).

Tissues with high energy requirements — such as the heart, liver, and brain — are particularly sensitive to mtDNA instability. Disruption of mtDNA ploidy in these tissues can trigger pathological cascades, contributing to the development of age-related diseases including Parkinson’s disease, Alzheimer’s disease, sarcopenia, and cardiomyopathies (Wallace, 2010; Schon et al., 2012). Furthermore, mtDNA depletion is a hallmark of several mitochondrial syndromes (e.g., Pearson syndrome, progressive external ophthalmoplegia) and is associated with acquired conditions such as type 2 diabetes, neurodegeneration, and cancer (Picard et al., 2014; Gorman et al., 2015; Vyas et al., 2016). Although mitochondrial DNA (mtDNA) depletion is a well-established hallmark of various mitochondrial disorders, an excessive increase in mtDNA copy number can also be pathological. Elevated mtDNA levels have been associated with a range of diseases, including cancer, where they may reflect or promote oxidative stress, genomic instability, and disrupted cellular metabolism. High mtDNA copy number has been observed in several tumor types, such as lymphoma and thyroid carcinoma, and is linked to poor prognosis in certain cancers (Hu et al., 2016; Alwehaidah et al., 2024). Additionally, excessive mtDNA replication may impair mitochondrial transcription, increase the burden of deletions, and disrupt oxidative phosphorylation, ultimately contributing to mitochondrial dysfunction and disease progression (Ylikallio et al., 2010; Abd Radzaket al., 2022).

mtDNA ploidy is not a static marker but a dynamic and regulated parameter, modulated by physiological and environmental stimuli. Cellular stressors such as oxidative stress, hypoxia, and exposure to mutagens can induce changes in mitochondrial biogenesis and mtDNA replication. These responses are orchestrated by signaling pathways involving AMPK, NRF1/2, and the master regulator PGC-1α (Scarpulla, 2008). In some contexts, increased ploidy may serve as a compensatory mechanism, buffering the effects of accumulating mtDNA mutations and preserving mitochondrial function until a critical heteroplasmy threshold is reached (Rajasimha et al., 2008). Conversely, chronic depletion of mtDNA can lead to mitochondrial dysfunction, impaired antioxidant defense, and increased vulnerability to damage (Suomalainen & Battersby, 2018).

Given its bacterial origin, limited DNA repair capacity, and proximity to ROS production sites, mtDNA is highly susceptible to damage by both endogenous and exogenous mutagens (Kazak et al., 2012). These agents can cause point mutations, large-scale deletions, and duplications, as well as mtDNA breaks, compromising the integrity of the mitochondrial genome (Yakes & Van Houten, 1997; Bratic & Larsson, 2013). Despite increasing interest in mitochondrial biology, the molecular mechanisms underlying changes in mtDNA copy number in response to mutagenic stress remain poorly understood.

Several studies have explored how environmental and chemical mutagens affect the ploidy of mitochondrial DNA (mtDNA), revealing that copy number alterations may serve as early indicators of genotoxic and oxidative stress. These changes are often interpreted as compensatory responses to mitochondrial damage or disruptions in mitochondrial biogenesis.

One of the earliest investigations linking chemical exposure to mtDNA content was conducted by Pavanello et al. (2013), who reported elevated mtDNA copy number in peripheral blood leukocytes of workers exposed to benzene. The increase was dose-dependent, with a 4% rise at ≤10 ppm and a 15% rise at >10 ppm, suggesting a systemic mitochondrial response to benzene-induced oxidative damage. Similarly, a population-based study by Carugno et al. (2012) in northern Italy found that even low-level occupational exposure to benzene — well below regulatory thresholds — was associated with significant increases in mtDNA content. These findings reinforce the notion that mitochondrial genome regulation is sensitive to chronic chemical exposure, even at sub-toxic levels.

Exposure to polycyclic aromatic hydrocarbons (PAHs) has also been linked to mtDNA copy number alterations. Choi et al. (2023) showed that 1-nitropyrene (1-NP), a PAH derivative, induces significant mtDNA amplification in human cells. The increase likely reflects cellular attempts to compensate for mtDNA damage incurred during 1-NP metabolism. Furthermore, a recent meta-analysis by Avilés-Ramírez et al. (2022) synthesized data from 22 studies involving over 6,000 individuals, showing consistent associations between exposure to heavy metals, PAHs, particulate matter, and cigarette smoke with altered mtDNA copy number. Although some heterogeneity was noted, the overall direction indicated a tendency toward mtDNA amplification, likely reflecting stress-induced mitochondrial proliferation.

A study by Mutlu et al. (2012) examined the effects of ochratoxin A and methanol on mtDNA in Drosophila. The researchers observed an increase in mtDNA copy number in response to mitochondrial DNA damage induced by these substances, highlighting the role of mtDNA copy number alterations as a response to chemical-induced mitochondrial stress.

Eom et al. (2011) investigated the potential of mtDNA copy number and hnRNP A2/B1 protein levels as biomarkers for direct benzene exposure. Their study found that both markers were significantly elevated in individuals exposed to benzene, suggesting their utility in monitoring benzene-induced mitochondrial and cellular alterations.

Ionizing radiation has been reported to alter mtDNA copy number in a cell-type – dependent manner. For example, Maguire et al. (2014) found that a single 2 Gy γ-ray exposure decreased mtDNA copy number in human bronchial epithelial (BEAS-2B) cells, whereas mtDNA increased in lung fibroblasts under the same conditions. In this study, mtDNA was measured 5 days post-irradiation, and BEAS-2B cells showed ∼20–40% reduction (to ≈60–80% of control) while HFL-1 fibroblasts showed an increase. In aged mice exposed to whole-body irradiation, only slight increases in mtDNA were observed in kidney and liver. These results suggest that ionizing radiation can deplete mtDNA in certain cell types (perhaps due to oxidative mtDNA damage or impaired replication), although other studies often report compensatory increases in mtDNA after irradiation Ionizing radiation is another potent genotoxic factor known to influence mtDNA content. Recent work by Seino et al. (2025) revealed that gamma irradiation in cell lines and mice led to increased mtDNA copy numbers, accompanied by reduced integrity of mitochondrial genomes. Interestingly, this effect was also observed in the offspring of irradiated animals, suggesting potential transgenerational inheritance of mtDNA instability.

Intercalating agents that block mtDNA replication also deplete mtDNA. Ethidium bromide (EtBr) is a classic mtDNA-depleting agent: Warren et al. (2017) treated primary rat striatal neurons and astrocytes with EtBr and found a dose-dependent decrease in mtDNA. Quantitative PCR showed that EtBr reduced mtDNA copy number by >50% in neuron-enriched cultures (where neurons are highly sensitive) but had little effect in astrocytes. Another recent method-development study (Novotny et al., 2023) used EtBr ± 2′,3′-dideoxycytidine to create human bronchial epithelial cells: one week of 50 ng/mL EtBr reduced mtDNA by ∼95%, and the combination with 25 μM ddC nearly eliminated mtDNA. These results confirm that EtBr (an intercalating mutagen) effectively lowers mtDNA copy number in proliferating mammalian cells.

Also, arsenic (a metalloid mutagen) causes mitochondrial dysfunction and mtDNA depletion in cultured cells: Partridge et al. (2007) exposed human–hamster hybrid A_L_ cells to arsenic, observing impaired oxidative phosphorylation that correlated with depletion in mtDNA copy number.

Collectively, these studies provide compelling evidence that mtDNA ploidy is a dynamic parameter modulated by environmental genotoxins. These insights support the relevance of mtDNA copy number not only as a biomarker of exposure but also as a potential contributor to disease processes linked to mitochondrial dysfunction.

Understanding how different mutagens affect mitochondrial DNA (mtDNA) copy number contributes to our knowledge of mitochondrial responses to genotoxic stress. These findings may help clarify the mechanisms underlying mitochondrial adaptation and impairment under chemical exposure. However, the existing body of research — conducted under varying experimental conditions and using diverse model systems — remains fragmented, limiting our ability to draw unified conclusions about the impact of mutagens on mitochondrial function.

The aim of this study is to analyze how established and potential exogenous mutagens — with well-characterized or partially understood mechanisms of action — affect the ploidy of the mitochondrial genome. This approach enables the identification of patterns in mitochondrial elimination under mutagenic stress. The investigation is conducted in human induced pluripotent stem cells (iPSCs), a powerful model for studying genome stability, cellular stress responses, and long-term consequences of genotoxic exposure (Zhang et al., 2013; Hämäläinen et al., 2015). Understanding how mutagens influence mtDNA ploidy in iPSCs will provide a foundation for further exploration of mitochondrial adaptation mechanisms in a controlled genomic background.

## Materials and Methods

Whole-genome sequencing (WGS) data were used to detect mutations and assess mitochondrial DNA (mtDNA) ploidy. Sequencing was performed using Illumina HiSeq and NovaSeq platforms, with a minimum coverage depth of 30× for the nuclear genome. Raw cram files from the work Kucab and colleagues in 2019 (Kucab et al., 2019). Reads were aligned to the human reference genome GRCh38, including the mitochondrial chromosome (chrM), using BWA-MEM (version 0.7.17 (r1188)).

Mitochondrial genome ploidy for each sample was estimated based on the ratio of mtDNA coverage normalized on length of mitochondrial genome to autosomes coverage (excliding sex chromosomes), normalized on their length (Wang et. al., 2017). This value was interpreted as the relative number of mtDNA copies per cell, normalized to the nuclear genome sequencing depth. This approach accounts for technical variation in sequencing depth, alignment efficiency, and library size, and is widely used in similar studies (Guo et al., 2013). We excluded from ther further analysis one BaP sample as outlier as well as gamma irradiation and SSR, that had no control information.

To ensure data reliability, additional metrics were calculated, including the coefficient of variation for mtDNA coverage across genomic regions and the coverage distribution profile along the mitochondrial genome. These analyses helped identify and exclude artifacts caused by local coverage dropouts or enrichment due to secondary DNA structures or ambiguous alignment of repetitive sequences. Statistical analyses were performed using R (version 4.3.2) in RStudio (version 2023.12.1+402) and Python (version 3.11.6) in PyCharm (version 2023.3.5). For data processing and visualization, we used the *ggplot2* (v3.4.4) package in R, and *pandas* (v2.1.4) and *matplotlib* (v3.8.2) in Python. All analyses were conducted in a controlled computing environment to ensure reproducibility.

## Results

To investigate the impact of mutagenic exposure on mitochondrial genome stability, we analyzed whole-genome sequencing (WGS) data from the study by Kucab et al. (2019), with permission from the original authors. The dataset includes human induced pluripotent stem cells (iPSCs) treated with 79 distinct mutagens representing a broad range of chemical classes and mechanisms of action. Each mutagen was applied at one to three different concentrations, depending on cytotoxicity and solubility constraints. In some cases, mutagens were administered in combination with an S9 metabolic activation mix, as their mutagenic potential is dependent on metabolic conversion. While the original study focused exclusively on mutations in the nuclear genome, mitochondrial DNA (mtDNA) was not examined. The dataset also includes several types of negative controls, such as untreated cells, water, culture medium, sodium chloride solution, and dimethyl sulfoxide (DMSO), depending on the solvent used for each mutagen.

Mitochondrial DNA (mtDNA) ploidy, calculated as the ratio of median coverage across the mitochondrial chromosome to the median coverage across autosomes, revealed significant alterations in mtDNA copy number following exposure to various mutagens. Among the 108 tested samples, 66 showed a statistically significant decrease in mtDNA ploidy compared to control samples (p < 0.05), whereas 9 mutagens induced an increase in mtDNA copy number (Fig. 1). The remaining compounds either did not cause statistically significant changes or exhibited high inter-replicate variability, preventing definitive conclusions.

**Fig. 1.**
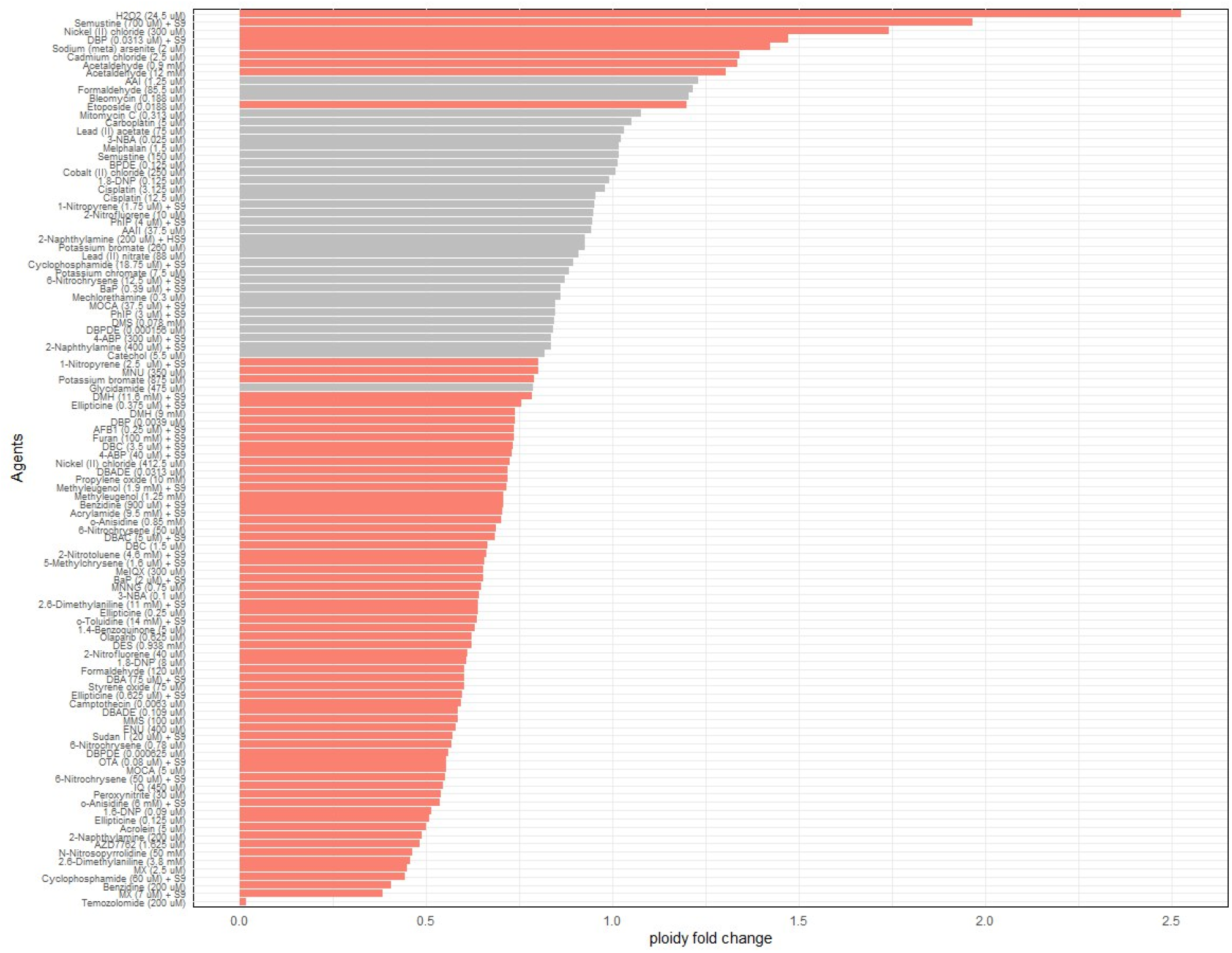
Ratio of median ploidy in mutagen-treated samples relative to control samples for each tested agent. Statistically significant differences compared to control are shown as pink bars; non-significant differences are shown in gray.

The most pronounced reductions in ploidy were observed after treatment with compounds known to impair DNA replication or induce oxidative damage, such as MX in samples 7 µM + S9 and 2.5 µM (Hyttinen et. al., 1995), benzidine (200 µM) (Phillips et.al., 1990) and cyclophosphamide (60 µM) + S9 (Gu et. al., 2023). These samples showed an average decrease in mtDNA copy number ranging from 25% to 60% relative to the corresponding control groups. For instance, cells treated with benzidine (200 µM) exhibited a median ploidy of 184, compared to 453 in controls (p = 1.24 × 10^−14^, Wilcoxon test), indicating substantial depletion of the mitochondrial pool due to disruption of the transcription-replication balance in mitochondria.

Conversely, samples treated with hydrogen peroxide (24.5 µM), semustine (700 µM) + S9, and nickel (II) chloride (300 µM) demonstrated statistically significant increases in mtDNA copy number (median increase of 30–50%, p < 0.01), likely reflecting a compensatory cellular response to stress and/or damage to individual mtDNA molecules. In some cases, an expansion in the range of ploidy values between replicates was also observed, potentially indicating unstable or cell-specific regulation of mitochondrial biogenesis in response to damage (Clay Montier et.al, 2009, Filograna et.al., 2021).

Under control conditions (untreated cells and solvent controls), mtDNA ploidy showed a stable distribution with a median value of approximately 379 copies per cell and relatively low dispersion (interquartile range: 355–453). Due to the non-normal distribution of values, non-parametric tests were used to assess statistical significance between control and treated groups: the Mann–Whitney (Wilcoxon) test and Fisher’s exact test for categorical analysis of ploidy increases or decreases.

Overall, 69% of the compounds caused statistically significant deviations from control values (p < 0.05, Benjamini–Hochberg corrected). Using Fisher’s test to compare the proportion of samples with mtDNA ploidy below a threshold of 200, significant enrichment was found among the groups treated with cyclophosphamide (60 µM) + S9 and N-nitrosopyrrolidine (50 mM) (p < 0.01). The rationale for adopting this cutoff is supported by previous studies demonstrating that mtDNA copy number is tightly regulated during differentiation and development to meet the energetic and biosynthetic demands of the cell (St John, 2014; Fukunaga, 2021). Furthermore, methodological studies have shown that robust quantitative PCR approaches consistently identify ∼200 copies as a critical point below which mitochondrial dysfunction becomes increasingly likely (Refinetti et al., 2017). Thus, using 200 copies as a threshold provides a practical and biologically relevant criterion for categorizing mtDNA content in the context of mitochondrial health and cellular homeostasis. Correlation analysis revealed a weak but statistically significant negative association between mtDNA ploidy and the number of detected point mutations (Spearman ρ = –0.31, p = 0.008), which may reflect impaired mtDNA replication and repair under conditions of mtDNA depletion.

## Discussion

The results of this study demonstrate that exposure to mutagenic agents in induced pluripotent stem cells (iPSCs) can cause statistically significant changes in mitochondrial DNA (mtDNA) ploidy, lead to the accumulation of specific somatic mutations, and, in some cases, result in rare structural rearrangements of the mitochondrial genome. The observed reduction in mtDNA copy number in certain treated samples indicates that specific chemical agents may disrupt mitochondrial replication and distribution processes or affect mitochondrial genome stability, potentially altering the functional state of the cells.

Notably, the majority of the tested compounds — despite their well-known mutagenic effects on sequence of the nuclear genome — did not induce significant changes in mitochondrial ploidy. This observation supports the hypothesis that specialized protective mechanisms exist to shield the mitochondrial genome from mutagenic stress. Potential factors contributing to this resilience include the physical segregation of mitochondria, which limits the diffusion of reactive metabolites into the organelles; the inherently high mtDNA copy number, providing a buffering capacity against partial damage; and the role of mitophagy — the selective removal of damaged mitochondria before they accumulate in the cell. In addition, although mitochondria lack nucleotide excision repair (NER), they retain a functioning base excision repair (BER) system capable of eliminating a broad range of mutagenic lesions, including oxidative and alkylation damage [Bohr et al., 2002; Gitschlag et al., 2020; Ashrafi & Schwarz, 2013].

Future research should focus on assessing the phenotypic consequences of these mitochondrial alterations. It is essential to determine how mtDNA depletion affects mitochondrial function, including ATP synthesis, reactive oxygen species (ROS) production, and the overall metabolic status of the cell. A key question is the temporal nature of the observed effects: are they stable and persistent, or reversible and reflective only of an acute cellular stress response? Clarifying cell- and tissue-specific differences in response to mutagenic exposure will help explain why mitochondrial disorders predominantly affect tissues with high energy demand and variable mitochondrial load.

Furthermore, based on the data presented, one may hypothesize the existence of relatively “mitochondria-sparing” mutagens that exert strong nuclear genotoxic effects with minimal mitochondrial impact. Such selective activity is of particular interest for the development of chemotherapeutic strategies with reduced mitochondrial toxicity.

In summary, these findings underscore the value of assessing mtDNA ploidy as a marker of cellular resilience to genotoxic stress and open new directions for exploring mitochondrial genome protection mechanisms. A deeper understanding of these processes holds promise for applications in both toxicology and regenerative medicine, where maintaining mitochondrial stability is becoming increasingly critical in the development of safe and effective cell-based therapies.

## Notes

### Competing Interest Statement

The authors have declared no competing interest.

### Summary of Updates

The authorship of the article has been corrected.

## References

1. Abd Radzak, S. M., Mohd Khair, S. Z. N., Ahmad, F., Patar, A., Idris, Z., & Yusoff, A. A. M. (2022). Insights regarding mitochondrial DNA copy number alterations in human cancer. International Journal of Molecular Medicine, 50(2), 104. doi: 10.3892/ijmm.2022.5160

2. Alwehaidah, M. S., Al-Awadhi, R., Roomy, M. A., & Baqer, T. A. (2024). Mitochondrial DNA copy number and risk of papillary thyroid carcinoma. BMC Endocrine Disorders, 24(1), 138. doi: 10.1186/s12902-024-01669-3

3. Anderson, S., Bankier, A.T., Barrell, B.G., de Bruijn, M.H., Coulson, A.R., Drouin, J., et al. Sequence and organization of the human mitochondrial genome. Nature. 1981 Apr;290(5806):457–465. doi: 10.1038/290457a0

4. Ashrafi, G., & Schwarz, T. L. (2013). The pathways of mitophagy for quality control and clearance of mitochondria. Cell Death & Differentiation, 20(1), 31–42. doi: 10.1038/cdd.2012.81

5. Avilés-Ramírez, C., Moreno-Godínez, M. E., Bonner, M. R., Parra-Rojas, I., Flores-Alfaro, E., Ramírez, M., Huerta-Beristain, G., & Ramírez-Vargas, M. A. (2022). Effects of exposure to environmental pollutants on mitochondrial DNA copy number: a meta-analysis. Environmental science and pollution research international, 29(29), 43588–43606. doi: 10.1007/s11356-022-19967-5

6. Bohr, V. A., Stevnsner, T., & Spelbrink, J. N. (2002). Mitochondrial DNA repair of oxidative damage in mammalian cells. Free Radical Biology and Medicine, 32(11), 1102– 1115. doi: 10.1016/S0891-5849(02)00878-6

7. Bratic, A., Larsson, N.G. The role of mitochondria in aging. J Clin Invest. 2013 Mar;123(3):951–957. doi: 10.1172/JCI64125

8. Carugno, M., Pesatori, A. C., Dioni, L., Hoxha, M., Bollati, V., Albetti, B., Byun, H. M., Bonzini, M., Fustinoni, S., Cocco, P., Satta, G., Zucca, M., Merlo, D. F., Cipolla, M., Bertazzi, P. A., & Baccarelli, A. (2012). Increased mitochondrial DNA copy number in occupations associated with low-dose benzene exposure. Environmental health perspectives, 120(2), 210–215. doi: 10.1289/ehp.1103979

9. Choi, S. H., Ochirpurev, B., Jo, H. Y., Won, J. U., Toriba, A., & Kim, H. (2023). Effects of polycyclic aromatic hydrocarbon exposure on mitochondrial DNA copy number. Human & experimental toxicology, 42, 9603271231216968. doi: 10.1177/09603271231216968

10. Clay Montier, L. L., Deng, J. J., & Bai, Y. (2009). Number matters: control of mammalian mitochondrial DNA copy number. Journal of genetics and genomics = Yi chuan xue bao, 36(3), 125–131. doi: 10.1016/S1673-8527(08)60099-5

11. Eom, H. Y., Kim, H. R., Kim, H. Y., Han, D. K., Baek, H. J., Lee, J. H., … & Shin, M. G. (2011). Mitochondrial DNA copy number and hnRNP A2/B1 protein: biomarkers for direct exposure of benzene. Environmental Toxicology and Chemistry, 30(12), 2762–2770. doi: 10.1002/etc.675

12. Filograna, R., Mennuni, M., Alsina, D., & Larsson, N. G. (2021). Mitochondrial DNA copy number in human disease: the more the better?. FEBS letters, 595(8), 976–1002. doi: 10.1002/1873-3468.14021

13. Finck, B. N., & Kelly, D. P. (2006). PGC-1 coactivators: inducible regulators of energy metabolism in health and disease. The Journal of clinical investigation, 116(3), 615–622. doi: 10.1172/JCI27794

14. Fukunaga, H. (2021). Mitochondrial DNA Copy Number and Developmental Origins of Health and Disease (DOHaD). International Journal of Molecular Sciences, 22(12), 6634. doi: 10.3390/ijms22126634

15. Gitschlag B.L., Kirby, C.S., Samuels, D.C., Gangula, R.D., Mallal, S.A., Patel, M.R. Homeostatic responses regulate selfish mitochondrial genome dynamics in C. elegans. Cell Metab. 2020 Dec 1;32(6):885–900.e6. doi: 10.1016/j.cmet.2020.10.001

16. Gorman, G.S., Schaefer, A.M., Ng, Y., Gomez, N., Blakely, E.L., Alston, C.L., et al. Prevalence of nuclear and mitochondrial DNA mutations related to adult mitochondrial disease. Ann Neurol. 2015 May;77(5):753–759. doi: 10.1002/ana.24362

17. Gu, L., Hickey, R. J., & Malkas, L. H. (2023). Therapeutic Targeting of DNA Replication Stress in Cancer. Genes, 14(7), 1346. doi: 10.3390/genes14071346

18. Guo, Y., Zhang, Q., Zhang, L., & Zhang, Y. (2013). A bioinformatics pipeline for estimating mitochondrial DNA copy number from whole-genome sequences. BMC Bioinformatics, 14, 1–7. doi: 10.1186/1471-2105-14-1

19. Hämäläinen, R.H., Manninen, T., Koivumäki, H., Kislin, M., Otonkoski, T., Suomalainen, A. Tissue- and cell-type–specific manifestations of heteroplasmic mtDNA 3243A>G mutation in human induced pluripotent stem cell–derived disease model. Cell Rep. 2015 Oct 20;13(3):647–660. doi: 10.1016/j.celrep.2015.09.025

20. Hu, L., Yao, X., & Shen, Y. (2016). Altered mitochondrial DNA copy number contributes to human cancer risk: evidence from an updated meta-analysis. Scientific reports, 6, 35859. doi: 10.1038/srep35859

21. Hyttinen, J. M., & Jansson, K. (1995). PM2 DNA damage induced by 3-chloro-4-(dichloromethyl)-5-hydroxy-2 (5H)-furanone (MX). Mutation research, 348(4), 183–186. doi: 10.1016/0165-7992(95)90007-1

22. Kazak, L., Reyes, A., Holt, I.J. Minimizing the damage: repair pathways keep mitochondrial DNA intact. Nat Rev Mol Cell Biol. 2012 Oct;13(10):659–671. doi: 10.1038/nrm3439

23. Kucab, J. E., Zou, X., Morganella, S., Joel, M., Nanda, A. S., Nagy, E., Gomez, C., Degasperi, A., Harris, R., Jackson, S. P., Arlt, V. M., Phillips, D. H., & Nik-Zainal, S. (2019). A Compendium of Mutational Signatures of Environmental Agents. Cell, 177(4), 821–836.e16. doi: 10.1016/j.cell.2019.03.001

24. Li, H., Handsaker, B., Wysoker, A., Fennell, T., Ruan, J., Homer, N., … & Durbin, R. (2009). The Sequence Alignment/Map format and SAMtools. Bioinformatics, 25(16), 2078–2079. doi: 10.1093/bioinformatics/btp352

25. Maguire, D., Zhang, S. B., & Okunieff, P. (2014). Mitochondrial genetic abnormalities after radiation exposure. Advances in experimental medicine and biology, 812, 1– 7. doi: 10.1007/978-1-4939-0620-8_1

26. Mutlu A. G. (2012). Increase in mitochondrial DNA copy number in response to ochratoxin A and methanol-induced mitochondrial DNA damage in Drosophila. Bulletin of environmental contamination and toxicology, 89(6), 1129–1132. doi: 10.1007/s00128-012-0826-1

27. Novotny, M. V., Xu, W., Mulya, A., Janocha, A. J., & Erzurum, S. C. (2024). Method for depletion of mitochondria DNA in human bronchial epithelial cells. MethodsX, 12, 102497. doi: 10.1371/journal.pone.0190456

28. Partridge, M. A., Huang, S. X., Hernandez-Rosa, E., Davidson, M. M., & Hei, T. K. (2007). Arsenic induced mitochondrial DNA damage and altered mitochondrial oxidative function: implications for genotoxic mechanisms in mammalian cells. Cancer research, 67(11), 5239–5247. doi: 10.1158/0008-5472.CAN-07-0074

29. Pavanello, S., Dioni, L., Hoxha, M., Fedeli, U., Mielzynska-Svach, D., & Baccarelli, A. A. (2013). Mitochondrial DNA copy number and exposure to polycyclic aromatic hydrocarbons. Cancer epidemiology, biomarkers & prevention: a publication of the American Association for Cancer Research, cosponsored by the American Society of Preventive Oncology, 22(10), 1722–1729. doi: 10.1158/1055-9965.EPI-13-0118

30. Phillips, D. H., Cross, M. F., Kennelly, J. C., Wilcox, P., & O’Donovan, M. R. (1990). Determination of benzidine--DNA adduct formation in CHO, HeLa, L5178Y, TK6 and V79 cells. Mutagenesis, 5 Suppl, 67–69. doi: 10.1093/mutage/5.Supplement.67

31. Picard, M., Zhang, J., Hancock, S., Derbeneva, O., Golhar, R., Golik, P., … & Wallace, D. C. (2014). Progressive increase in mtDNA 3243A> G heteroplasmy causes abrupt transcriptional reprogramming. Proceedings of the National Academy of Sciences, 111(38), E4033–E4042. doi: 10.1073/pnas.1414028111

32. Rajasimha, H.K., Chinnery PF, Samuels DC. Selection against pathogenic mtDNA mutations in a stem cell population leads to the loss of the 3243A->G mutation in blood. Am J Hum Genet. 2008 Feb;82(2):333–343. doi: 10.1016/j.ajhg.2007.10.005

33. Refinetti, P., Warren, D., Morgenthaler, S., & Ekstrøm, P. O. (2017). Quantifying mitochondrial DNA copy number using robust regression to interpret real time PCR results. BMC research notes, 10, 1–7. doi: 10.1186/s13104-017-2913-1

34. Robin, E.D., Wong, R. Mitochondrial DNA molecules and virtual number of mitochondria per cell in mammalian cells. J Cell Physiol. 1988 Sep;136(3):507–513. doi:10.1002/jcp.1041360316

35. Scarpulla, R.C. Nuclear control of respiratory gene expression in mammalian cells. J Cell Biol. 2008 Apr 7;181(1):9–16. doi: 10.1083/jcb.200801040

36. Schon, E.A., Di Mauro, S., Hirano, M. Human mitochondrial DNA: roles of inherited and somatic mutations. Nat Rev Genet. 2012 Dec;13(12):878–890. doi: 10.1038/nrg3275

37. Seino, R., Kubo, H., Nishikubo, K., & Fukunaga, H. (2025). Radiation-induced impacts on mitochondrial DNA and the transgenerational genomic instability. Environment international, 196, 109315. doi: 10.1016/j.envint.2025.109315

38. Spinelli, J.B., Haigis, M.C. The multifaceted contributions of mitochondria to cellular metabolism. Nat Cell Biol. 2018 Jul;20(7):745–754. doi: 10.1038/s41556-018-0124-1

39. St John J. (2014). The control of mtDNA replication during differentiation and development. Biochimica et biophysica acta, 1840(4), 1345–1354. doi: 10.1016/j.bbagen.2013.10.036

40. Suomalainen, A., Battersby, B.J. Mitochondrial diseases: the contribution of organelle stress responses to pathology. Nat Rev Mol Cell Biol. 2018 Feb;19(2):77–92. doi: 10.1038/nrm.2017.66

41. Vyas, S., Zaganjor, E., Haigis, M.C. Mitochondria and cancer. Cell. 2016 Jul 28;166(3):555–566. doi: 10.1016/j.cell.2016.07.002

42. Wai, T., Teoli, D., Shoubridge, E.A. The mitochondrial DNA genetic bottleneck results from replication of a subpopulation of genomes. Nat Genet. 2008 Dec;40(12):1484–1488. doi: 10.1038/ng.258

43. Wallace, D.C. Bioenergetics in human evolution and disease: implications for the origins of biological complexity and the missing genetic variation of common diseases. Philos Trans R Soc Lond B Biol Sci. 2010 Jan 27;365(1544):341–350. doi: 10.1098/rstb.2009.0198

44. Wang, W., Liu, W. M., Yang, C., Chou, C. H., Chen, H. J., Wu, H. M., & Lee, R. K. K. (2017). Mitochondrial DNA content is associated with ploidy status, maternal age, and oocyte maturation methods in mouse blastocysts. International Journal of Molecular Sciences, 18(5), 979. doi: 10.3390/ijms18050979

45. Warren, E.B., Aicher, A.E., Fessel, J.P., Konradi, C. (2017) Mitochondrial DNA depletion by ethidium bromide decreases neuronal mitochondrial creatine kinase: Implications for striatal energy metabolism. PLOS ONE 12(12): e0190456. doi: 10.1371/journal.pone.0190456

46. Yakes, F.M., Van Houten, B. Mitochondrial DNA damage is more extensive and persists longer than nuclear DNA damage in human cells following oxidative stress. Proc Natl Acad Sci U S A. 1997 Jan 21;94(2):514–519. doi: 10.1073/pnas.94.2.514

47. Ylikallio, E., Tyynismaa, H., Tsutsui, H., Ide, T., & Suomalainen, A. (2010). High mitochondrial DNA copy number has detrimental effects in mice. Human molecular genetics, 19(13), 2695–2705. doi: 10.1093/hmg/ddq163

48. Zhang, Y., Pak, C., Han, Y., Ahlenius, H., Zhang, Z., Chanda, S., et al. Rapid single-step induction of functional neurons from human pluripotent stem cells. Neuron. 2013 Jun 5;78(5):785–798. doi: 10.1016/j.neuron.2013.05.029

